# SERINC5 co-expressed with HIV-1 Env or present in a target membrane destabilizes small fusion pores leading to their collapse

**DOI:** 10.1101/2025.09.09.675038

**Authors:** Ruben M. Markosyan, Mariana Marin, Gregory B. Melikyan

## Abstract

SERINC5 is a multipass membrane protein that reduces infectivity of retroviruses and other enveloped viruses through incorporation into budding virions and inhibiting viral fusion. Several modes of anti-HIV activity of SERINC5 have been reported, including binding to Env glycoprotein, induction of conformational changes and destabilization of Env, as well as disruption of trans-membrane asymmetry of viral envelope. All these reported anti-HIV mechanisms involve SERINC5 incorporation into progeny virions, while there is very little information regarding a potential antiviral activity of SERINC5 in target cells. Here, we show that SERINC5 expressed in target cells efficiently inhibits fusion with cells expressing sensitive but not resistant HIV-1 strains. We show that this activity does not result from downregulation of CD4 or coreceptors on target cells or interference with the formation of ternary Env/CD4/coreceptor complexes. We further demonstrate that SERINC5 destabilizes small fusion pores causing their collapse and that this block can be rescued by incorporation of phosphatidylserine to either effector (Env-expressing) or target cells. Interestingly, we did not detect a significant reduction of HIV-1 pseudovirus fusion with SERINC5 expressing target cells, while virus-mediated cell-cell fusion (fusion-from-without) was inhibited, suggesting a potential role of virus’ entry pathway in SERINC5 restriction. Collectively, our results reveal a novel mechanism of inhibition of HIV-1 fusion by SERINC5 through destabilization of small fusion pores in a manner that depends on lipid composition.

**Importance:** SERINC5 incorporates into virions produced by infected cells and inhibits viral fusion with target cells through a poorly understood mechanism. Here, we show that SERINC5 blocks HIV-1 Env mediated cell fusion when expressed in either effector (Env-expressing) or target cells. Inhibition of Env-mediated fusion by SERINC5 expressed in target cells is not through reduction in receptor or coreceptor expression or interference with Env’s ability to engage a requisite number of receptors and coreceptors. We demonstrate that the block of fusion is at a post-hemifusion stage of small fusion pores that collapse when SERINC5 is present in effector or target membrane and that this block is rescued by incorporation of specific anionic lipids, such as phosphatidylserine. These findings reveal a previously unappreciated mode of HIV-1 restriction through destabilization of small fusion pores that occurs irrespective of SERINC5 localization in either of fusing membranes.

## Introduction

Serine incorporator proteins 3 and 5 (SERINC3 and SERINC5) are 10-transmembrane domain proteins that reduce retrovirus infectivity by incorporating into virions and diminishing their ability to fuse with target cells (1–5). While SERINC expression is not increased upon interferon stimulation, endogenous levels of these proteins in CD4 T cells restrict HIV-1 infection (3, 5). The cellular functions of SERINC family proteins are not well understood (6, 7), but SERINC5 has been recently shown to exhibit a lipid scrambling activity manifested in phosphatidylserine (PS) exposure on the virion surface (8, 9). Interestingly, recent studies documented a broad antiviral activity of SERINC5 (hereafter abbreviated SER5) against diverse enveloped viruses, including the Influenza A virus (IAV), SARS-CoV-2, Classical Swine Fever virus, and others (10–14). Viruses counteract SER5 restriction by encoding proteins that bind to and degrade and/or remove SER5 from plasma membrane, thereby preventing its incorporating into virions. These include HIV-1 Nef, Murine Leukemia virus GlycoGag, Equine Infectious Anemia virus S2, and SARS-CoV-2 ORF7a proteins (3, 5, 10, 15–17).

In the absence of Nef, HIV-1’s resistance to SER5 restriction maps to Env glycoprotein, with lab-adapted Tier 1 strains being sensitive and primary isolates being relatively resistant to this protein (1, 3–5, 10, 18–20). The gp120 variable loop 3 (V3), which controls virus coreceptor tropism and Env stability (21), has been identified as the major determinant of SER5 resistance (1). The gp41 cytoplasmic tail has also been implicated to Env’s sensitivity to SER5 (22), but this notion has not been confirmed by other groups, including ours (4, 23). The mechanism of SER5’s antiviral activity and virus resistance is not well-understood and appears to be multifaceted. SER5 has been shown to alter the conformation of Env on virions, promote its spontaneous inactivation and sensitize to neutralizing antibodies (1, 4, 18, 19, 24, 25). These SER5 effects have been proposed to occur through direct SER5-Env interaction that is promoted upon Env-CD4 engagement (20). However, our 2-color super-resolution microscopy results do not support direct interaction of Env and SER5 on single virions (26), while revealing SER5-mediated disruption of Env clusters on mature HIV-1 that may diminish virus’ fusion competence. A recent study has linked the lipid scrambling activity of SER5 to reduction of HIV-1 infectivity (8), although it remains unclear whether loss of viral membrane’s lipid asymmetry strictly correlates with reduction in infectivity (9). Regardless of the exact mechanism of HIV-1 restriction, the antiviral activity of SER5 against other enveloped viruses is indicative of a universal mechanism of inhibition of viral fusion, perhaps through modification of lipid membranes, as has been proposed in (27).

Studies of SER5-mediated restriction have focused on the effects of this protein incorporated into viral membrane. Only two groups have examined the anti-IAV activity of SER5 expressed in target cells but arrived at discordant conclusions regarding its ability to inhibit IAV HA-mediated fusion/infection (13, 14). It is thus unclear whether SER5 can inhibit viral fusion when present in target cells, which could prompt a revision of the current models of SER5-mediated virus restriction. Here, we conducted a series of experiments to assess the effects of SER5 expressed in effector (fusion protein-expressing) and target cells on HIV-1 Env-mediated infection and membrane fusion. These experiments revealed a marked inhibition of cell-cell fusion, but not virus-cell fusion or infection by SER5 in target cells. Inhibition of cell-cell fusion was not associated with downregulation of CD4 or coreceptor levels or with inhibition of Env binding to CD4 or coreceptors. We also found that SER5 in target cells diminished cell-cell fusion mediated by Env expressed on effector cells or Env incorporated into virions. Notably, the sensitivity of HIV-1 strains to SER5 restriction from effector cells was mirrored by SER5 expressed in target cells. Importantly, Env-mediated cell-cell fusion was arrested at a small pore stage, and addition of exogenous PS antagonized the SER5 effect on cell fusion. These results demonstrate the ability of SER5 to interfere with HIV-1 Env-mediated fusion when present in a target cell, suggesting a novel mechanism of antiviral activity.

## Results

### SER5 expressed both in effector and target cells inhibits cell-cell fusion mediated by sensitive HIV-1 Env

Using a dual-split protein assay (28), we have previously reported inhibition of HIV-1 Env-mediated cell-cell fusion by SER5 expressed in effector cells (4). Here, we confirmed the restriction activity of SER5 transiently expressed in the effector HeLa cells, using a fluorescence microscopy-based assay that detects redistribution of small cytosolic dyes between fused cells (29, 30) (Suppl. Fig. S1). As expected, co-expression of Nef rescued fusion with SER5-expressing effector cells. To test if SER5 can inhibit HIV-1 fusion when present in the target cells, we compared fusion of TF228.1.16 effector cell line stably expressing a CXCR4-tropic HIV-1 Env (31) to JTag CD4 T-cells endogenously expressing SER3 and SER5 and double-knockout (S3/S5−/−) JTag cells (5). To improve HIV-1 fusion efficiency these cells were additionally transduced with a CD4 expressing vector. Env-expressing TF228.1.16 cells fused with JTag depleted of SER3 and SER5 significantly better than with parental cells (Fig. 1A). Thus, endogenous levels of SER3/SER5 in JTag cells suppress HIV-1 Env-mediated cell fusion. Conversely, overexpression of SER3 or SER5 in target TZM-bl cells inhibited Env-mediated fusion with the TF.228.1.16 effector cells (Fig. 1B). By contrast, expression of the inactive member of SERINC family proteins, SER2 (4, 19), in target cells was without effect on cell fusion (Fig. 1B). The inhibition of cell-cell fusion by ectopically expressed SER5, but not SER2, observed in a microscopy-based assay was confirmed using a dual-split-GFP/Luciferase cell-cell fusion assay (Suppl. Fig. S2). Note that, this assay detected a slight reduction in cell-cell fusion upon SER2 expression in effector or target cells, which was, however, far less potent than expression of SER5.

**Figure 1.**
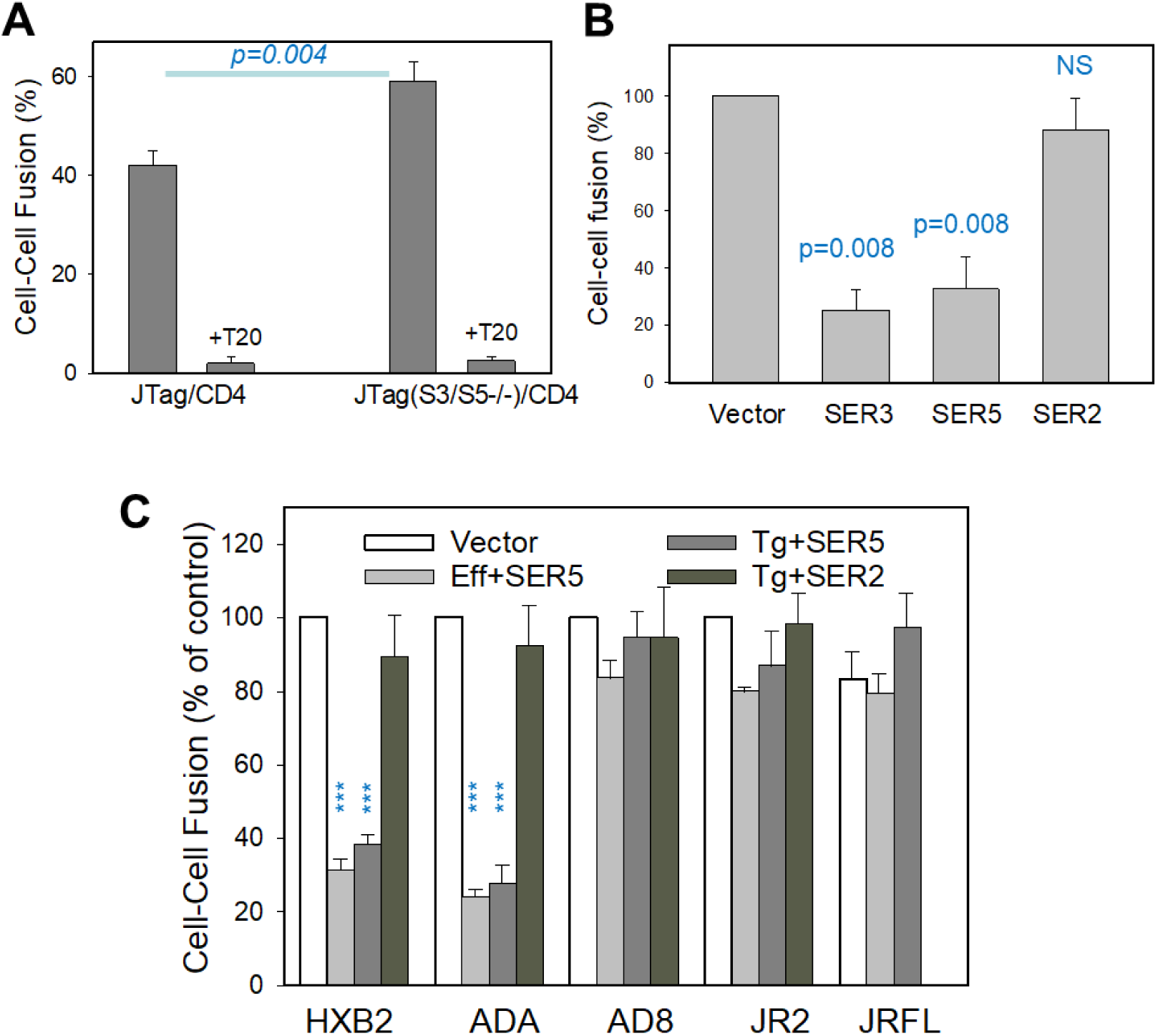
SER3 and SER5 in target cells inhibit HIV-1 Env-mediated fusion. (A) Parental JTag cells or JTag cells depleted of SER3 and SER5 (S3/S5−/−) were co-cultured with the TF228.1.16 effector cells stably expressing HIV-1 Env for 2 hours at 37 °C. To improve the fusion efficiency, JTag cells were transduced with a CD4-expressing retroviral vector. In control experiments, Env-mediated fusion was blocked by the inhibitory T20 peptide (40 nM). Cell-cell fusion was quantified using a microscopy-based assay. Mean and standard deviation from 3 independent experiments, each performed in duplicate, are shown. Statistical analysis was performed using Student’s t-test. (B) TF2281.16 cells were fused with TZM-bl target cells transfected or not with SER2, SER3 or SER5 plasmids. Cells were co-incubated for 2.5 hours at room temperature and then shifted to 37 °C for 30 minutes to allow fusion. Mean and standard deviation from 5 independent experiments, each performed in duplicate, are shown. Statistical analysis was done using Mann-Whitney test. (C) Expression of SER5 selectively reduces fusion of sensitive HXB2 (X4-tropic) and ADA (R5-tropic)), but not resistant R5-tropic (JRFL, JR2 and AD8) HIV-1 Env strains, irrespective to whether SER5 was expressed in target (Tg) TZM-bl or effector (Eff) HEK293T cells transiently expressing HIV-1 Env. In control experiments, SER2 had no measurable effect on fusion of sensitive or resistant Env strains. Fusion was assessed after 3 hours of incubation at 37 °C. Data are presented as means ± SEM from three (A), five (B), and three (C) independent experiments, each performed in duplicate. (*, p ≤ 0.05, **, p ≤ 0.01, ***, p ≤ 0.001, NS is not significant, using Student’s t-test).

The HIV-1 sensitivity to virion-incorporated SER5 maps to Env glycoprotein, specifically, to the gp120 V3 loop (1). We, therefore, asked if the sensitivity of Env-mediated fusion to SER5 expressed in target cells exhibits the same HIV-1 strain-dependence as when SER5 is expressed in effector cells. HEK293T cells transiently expressing Env (and SER5, when indicated) were co-cultured with TZM-bl cells transfected with SER5 or SER2 or mock-transfected. The resulting cell-cell fusion driven by HXB2 (lab-adapted, CXCR4-tropic) or ADA (lab-adapted, CCR5-tropic) Env was equally efficiently suppressed by SER5 in effector or target cells (Fig. 1C). By contrast, fusion mediated by JRFL, JR2 and AD8 Env glycoproteins that are relatively resistant to SER5 restriction (1, 3–5, 10, 18–20) was unaffected by SER5 expression in effector or target cells (Fig. 1C). In control experiments, SER2 failed to inhibit cell fusion mediated by any of these Env glycoproteins. Unless stated otherwise, all subsequent experiments were performed using HXB2 Env chosen for its high sensitivity to SER5-mediated restriction ((4, 18) and Figs. 1C and 2A).

**Figure 2.**
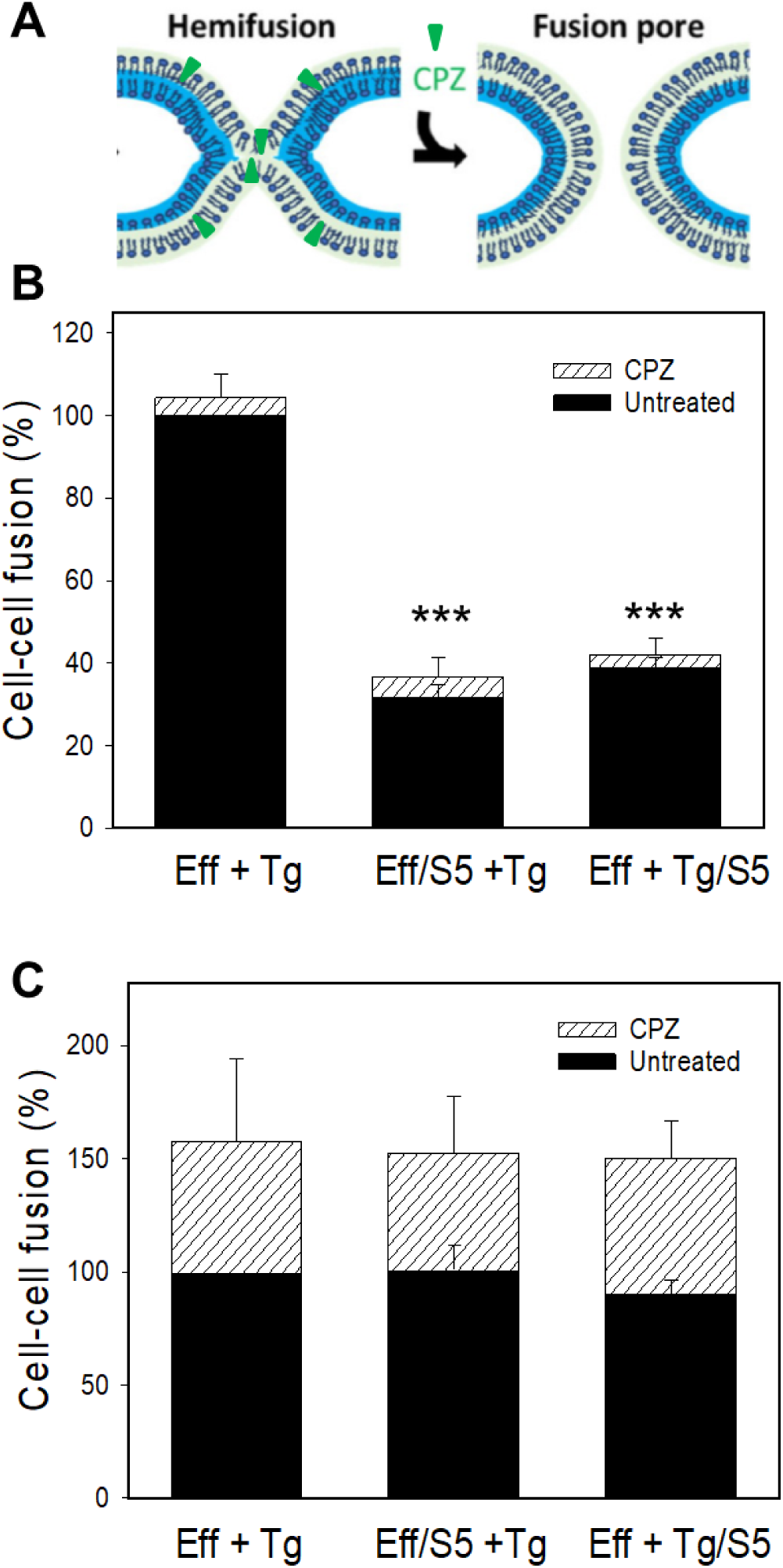
Inhibition of Env-mediated fusion by SER5 does not trap viral fusion at a hemifusion intermediate. (A) HEK293T effector cells transfected with HIV-1 HXB2 Env were fused with TZM-bl target cells. SER5 was expressed on either effector or target cells. Subsequent exposure to 0.5 mM CPZ for 1 min caused only a marginal increase in cell-cell fusion (hatched bars). (B) Same is in A, but the effector cells were transfected with influenza HA (X:31 strain) and pretreated with trypsin (10 µg/ml) and neuraminidase (0.1 mg/ml) for 5 min to activate HA. Cell co-cultures were exposed to pH 4.8 buffer for 10 min at room temperature and further incubated for 1 hour at 37 °C. No inhibition of fusion was observed with SERINC5 expression on either side. Exposure to 0.5 mM CPZ significantly enhanced HA-mediated fusion. Results are presented as mean ± SEM from three independent experiments, each performed in duplicate. Statistical analysis was performed using Student’s t-test.

Next, we asked if SER5 in target cells was active against non-HIV fusion proteins. Unlike the HIV-1 Env-mediated cell fusion (Fig. 2B), the Influenza virus X:31 HA-mediated fusion triggered by exposure to low pH was not inhibited by SER5 expressed in effector or target cells (Fig. 2C). The resistance of HA-mediated cell fusion to SER5 was also observed for another IAV HA strain, PR8/34 (Suppl. Fig. S3). Thus, SER5 selectively inhibits cell-cell fusion mediated by sensitive HIV-1 Env strains and not strains known to be resistant to restriction or IAV HA-mediated fusion.

### SER5 expression in target cells does not downregulate CD4/coreceptors or interfere with Env/CD4/coreceptor interactions

To determine if SER5 in target cells inhibits HIV-1 Env-mediated fusion by downregulating CD4 or coreceptor expression, we compared the levels of these proteins on control and SER5 expressing target cells. Immunofluorescence labeling for CD4 and CXCR4 and flow cytometry analysis did not reveal significant changes in surface expression of these proteins in control and SER5- or SER2-transfected TZM-bl cells (Suppl. Fig. S4). Further, overexpression of CD4 or coreceptors (CXCR4 or CCR5) did not rescue HXB2 or ADA Env-mediated fusion with SER5 expressing target cells (Suppl. Fig. S5). These results rule out downregulation of CD4 or coreceptor as the mechanism for inhibition of Env-mediated fusion by SER5 expressed in target cells.

Another potential mode of SER5 mediated inhibition of HIV-1 fusion is through interference with the binding of Env to CD4 and/or coreceptors on the cell surface, perhaps through restriction of their lateral mobility. To explore this possibility, we took advantage of the ability to capture/stabilize HIV-1 fusion at a step when a fraction of Env glycoproteins engages a requisite number of CD4 and coreceptors. We have demonstrated that preincubation of effector and target cells at a sub-threshold temperature for HIV-1 fusion (typically, 23 °C) allows Env to form ternary complexes with CD4 and cognate coreceptors but not to proceed to fusion (32, 33). The formation of Env/CD4/coreceptor complexes at this temperature-arrested stage (TAS) is manifested by acquisition of resistance to inhibitors of CD4 and CXCR4 binding (32, 33). As expected, prolonged pre-incubation of HXB2 Env-expressing effector cells and target cells at 23 °C rendered the subsequent fusion partially resistant to a fully inhibitory concentration of AMD3100 (CXCR4 binding inhibitor) added after preincubation (Fig. 3A). In contrast, the gp41-derived T20 peptide that prevents the formation of the final 6-helix bundle structure driving the formation of a fusion pore (32–34) abrogated fusion equally potently when present throughout the experiment or added at TAS (Fig. 3A). Like Env-mediated fusion with control cells, fusion with SER5-exprssing target cells arrested at TAS was partially resistant to AMD3100 (Fig. 3B). In fact, protection of fusion from AMD3100 added at TAS was independent of SER5 expression, with 61% and 67% of fusion of untreated cells occurring in the presence of AMD3100 for control and SER5-expressing cells, respectively (Fig. 3A, B). These results imply that SER5 does not interfere with the engagement of CD4 or CXCR4 by Env on the cell surface, at least on the time scale of establishing TAS.

**Figure 3.**
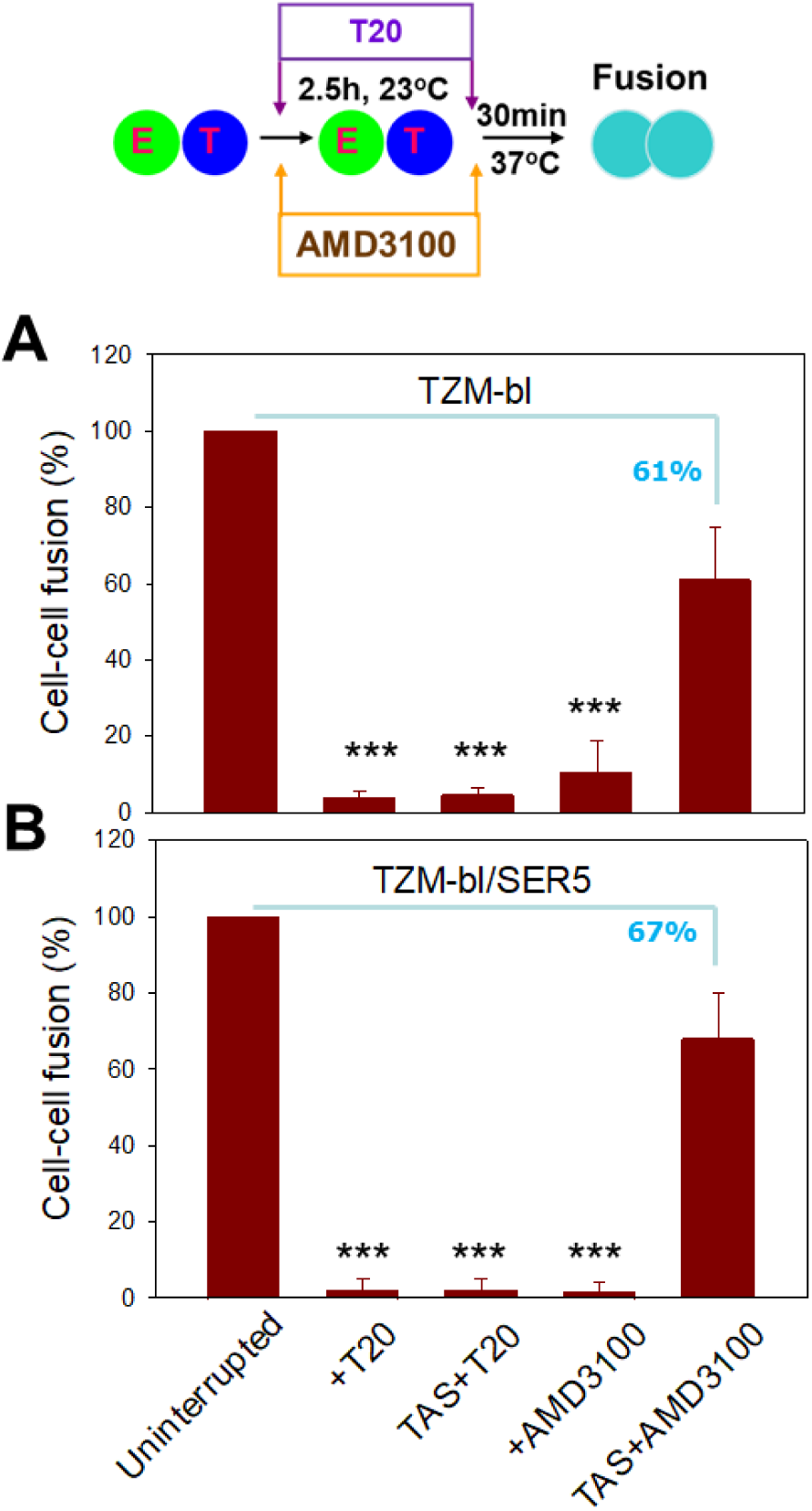
SER5 does not interfere with functional engagement of CD4 and CXCR4 by HIV-1 Env. HIV-1 Env-mediated cell fusion between effector HEK293T cells and TZM-bl cells was arrested by co-incubation at 23 °C for 2.5 hours, which enables engagement of CD4 and coreceptors but not fusion (referred to as a temperature-arrested stage, TAS). HEK293T cells were transfected with either HXB2 Env alone (A) or with HXB2 Env and SER5 (B). The CXCR4 binding inhibitor, AMD3100 (30 µM), or the gp41 6-helix bundle formation inhibitor, T20 (40 nM) were either present throughout the experiment or added at TAS, after preincubation at 23 °C. Following pre-incubation, cells were then shifted to 37 °C for 30 minutes to allow fusion. Mean and SEM are from four independent experiments, with each experiment performed in duplicate. Statistical analysis was performed using Student’s t-test.

### SER5 does not capture HIV-1 Env-mediated fusion at a hemifusion stage

Viral protein-mediated fusion proceeds through a hemifusion intermediate (e.g., (35–46)). Depending on the viral glycoprotein and experimental conditions, a significant fraction of fusion events may not progress beyond a hemifusion stage (36–39). These dead-end hemifusion structures can be converted to full fusion by a brief treatment with chlorpromazine (CPZ), which we have shown to partition into and destabilize the hemifusion diaphragm (Fig. 2A), thereby promoting fusion pore formation (38). In agreement with our previous findings (47–49), a sizeable fraction of HA-mediated cell fusion remained at a hemifusion stage, as evidenced by a nearly 30% increase in cell fusion after CPZ treatment (Fig. 2C, hatched bars). By comparison, HIV-1 Env is less prone to form dead-end hemifusion structures, as judged by a marginal increase in cell-cell fusion after exposure to CPZ (Fig. 2B). Importantly, the lack of rescue of Env-mediated fusion with SER5-expressing cells by CPZ, implies that SER5 does not arrest HIV-1 Env-mediated cell fusion at a hemifusion stage and is, thus, likely to block downstream steps of fusion.

### Phosphatidylserine and phosphatidylglycerol selectively rescue HIV-1 Env-mediated fusion of SER5-expressing cells

Since the lipid composition modulates viral protein-mediated fusion, and HIV-1 Env-mediated fusion, in particular (32, 50, 51), we tested whether the SER5-imposed block of Env-mediated fusion was lipid-dependent. An additional motivation was the recently discovered lipid scrambling activity of SER5, which has been proposed to contribute to the antiviral activity of this protein (8). Co-cultures of effector and target cells were pretreated with exogenously added phosphatidylcholine (PC), phosphatidylserine (PS), phosphatidylglycerol (PG) or bis(monoacylglycero)phosphate (BMP), and the resulting cell-cell fusion was measured after further incubation at 37 °C. Env-mediated fusion with control and SER2 expressing target cells was modestly promoted by exogenous PS and, to a lesser extent, PG (Fig. 4A). In contrast, PS and PG potently restored fusion with SER5 expressing cells almost to the level of SER5-negative cells (Fig. 4A). This selective antagonism of PS and PG with SER5-mediated inhibition of fusion can be readily appreciated by plotting the fold-inhibition of cell-cell fusion by SER5 in cells pretreated with different lipids (Fig. 4B). Thus, PS and PG specifically rescue fusion with SER5-expressing cells, whereas PC and BMP are without effect. The same lipid-dependence was observed upon SER5 co-expression with Env in effector cells. Here too, PS, but not PC or BMP, rescued fusion between SER5-expressing effector cells with target cells, while PC or BMP did not significantly modulate Env-mediated fusion (Fig. 4C, D).

**Figure 4.**
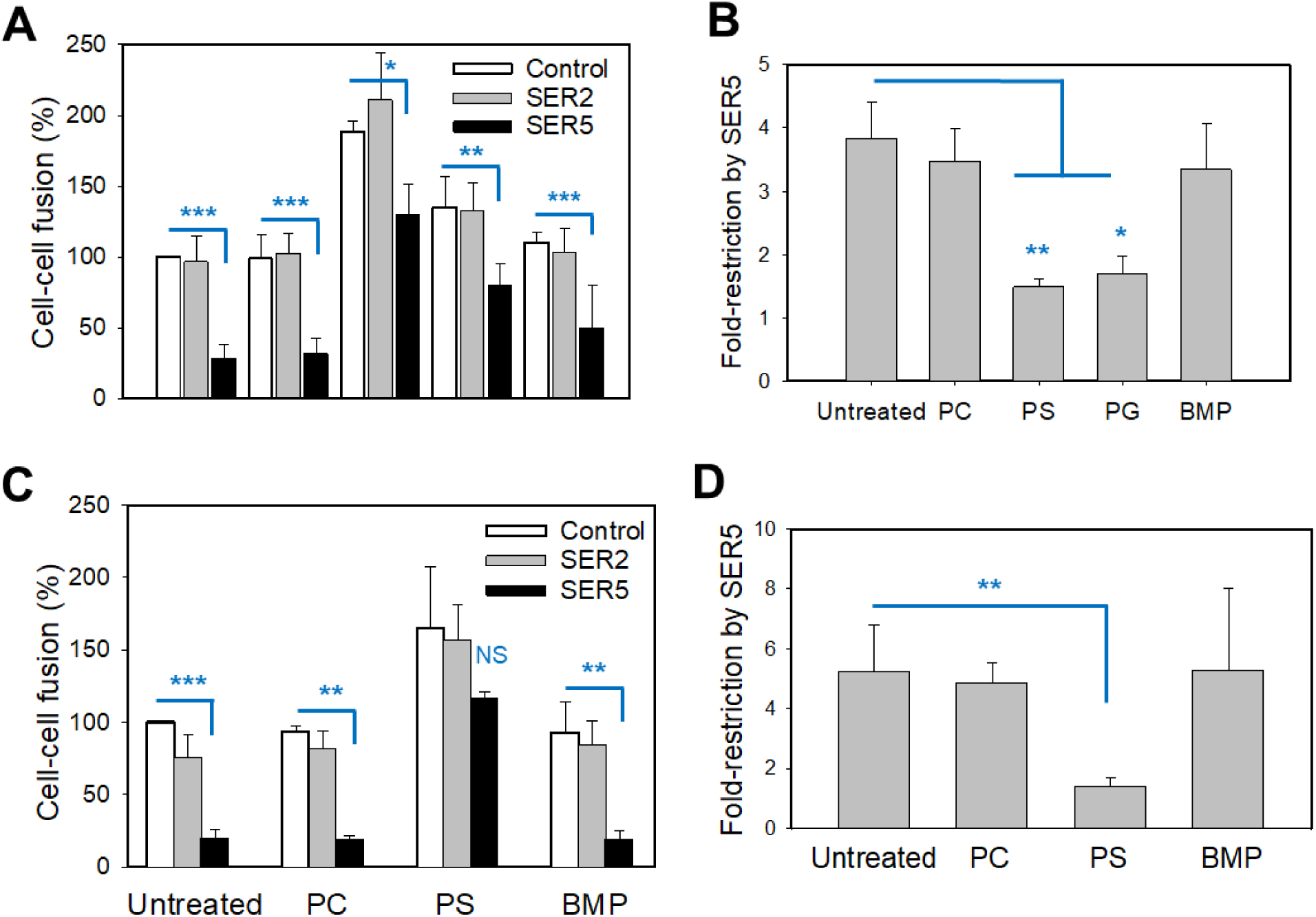
Lipid composition of the plasma membrane modulates SER5-mediated restriction of HIV-1 Env-mediated cell fusion. (A, B) TZM-bl target cells transiently expressing SER2 (negative control) or SER5 were co-incubated with HEK293T effector cells expressing HXB2 Env for 30-min at 23 °C. Cells were then treated with BSA (1mg/ml) containing freshly suspended lipids 10 µg/ml DOPC, DOPS, DOPG, or BMP for 20 min, washed twice with BSA-containing buffer to remove unincorporated lipids and incubated for 2.5 hours at 37 °C to allow fusion. The effect of exogenous lipids on the fold-restriction of cell fusion by SER5 is plotted in panel B. (C, D) Lipid-dependence of SER5-mediated restriction of HXB2 Env-mediated cell fusion between effector HEK293T cells ectopically expressing SER2 and SER5 with TZM-bl target cells. The effect of exogenous lipids on the fold-restriction of cell fusion by SER5 is plotted in panel D. Each value reflects the mean ± SEM from four (A) and two (C) independently repeated experiments, each performed in duplicate. Statistical analysis was performed using Student’s t-test.

We next asked if selective enhancement of fusion with SER5-expressing cells occurs through incorporation of PS in effector *vs* target cells. Target or effector cells were separately pretreated with exogenous lipids followed by removal of excess lipids and co-culture to allow fusion. Addition of PS selectively promoted fusion with SER5-expressing cells, irrespective of whether this lipid was added to effector or target cells (Fig. S6). Such a “symmetric” enhancing effect of PS suggests that late stages of Env-mediated fusion with SER5-expressing cells, perhaps downstream the membrane merger, are lipid-dependent. We hypothesized that SER5 and PS can modulate the stability of small fusion pores.

### SER5 slows down the formation of fusion pores and prevents their dilation

To resolve the formation of fusion pores in real time, we synchronized Env-mediated fusion by first capturing this process at TAS (preincubation at 23 °C) and quickly shifting to 37 °C using a custom-made temperature jump setup ((29, 52) and Fig. 5A). Owing to the formation of functional Env/CD4/coreceptor complexes, fusion from TAS is much faster than uninterrupted cell-cell fusion at 37 °C. Effector and target cells were co-cultured at 23 °C for 2.5 h. The formation of nascent fusion pores after shifting to 37 °C resulted in redistribution of calcein loaded into effector cells to target cells (Fig. 5C). These experiments revealed that the kinetic of fusion pore formation with SER5-expressing target cells captured at TAS was markedly slower than with control cells (Fig. 5B). This result, together with the observation that SER5 does not interfere with the formation of Env/CD4/CXCR4 complexes at TAS (Fig. 3), supports the notion that SER5 interferes with the late steps of Env-mediated membrane fusion, namely the formation of fusion pores.

**Figure 5.**
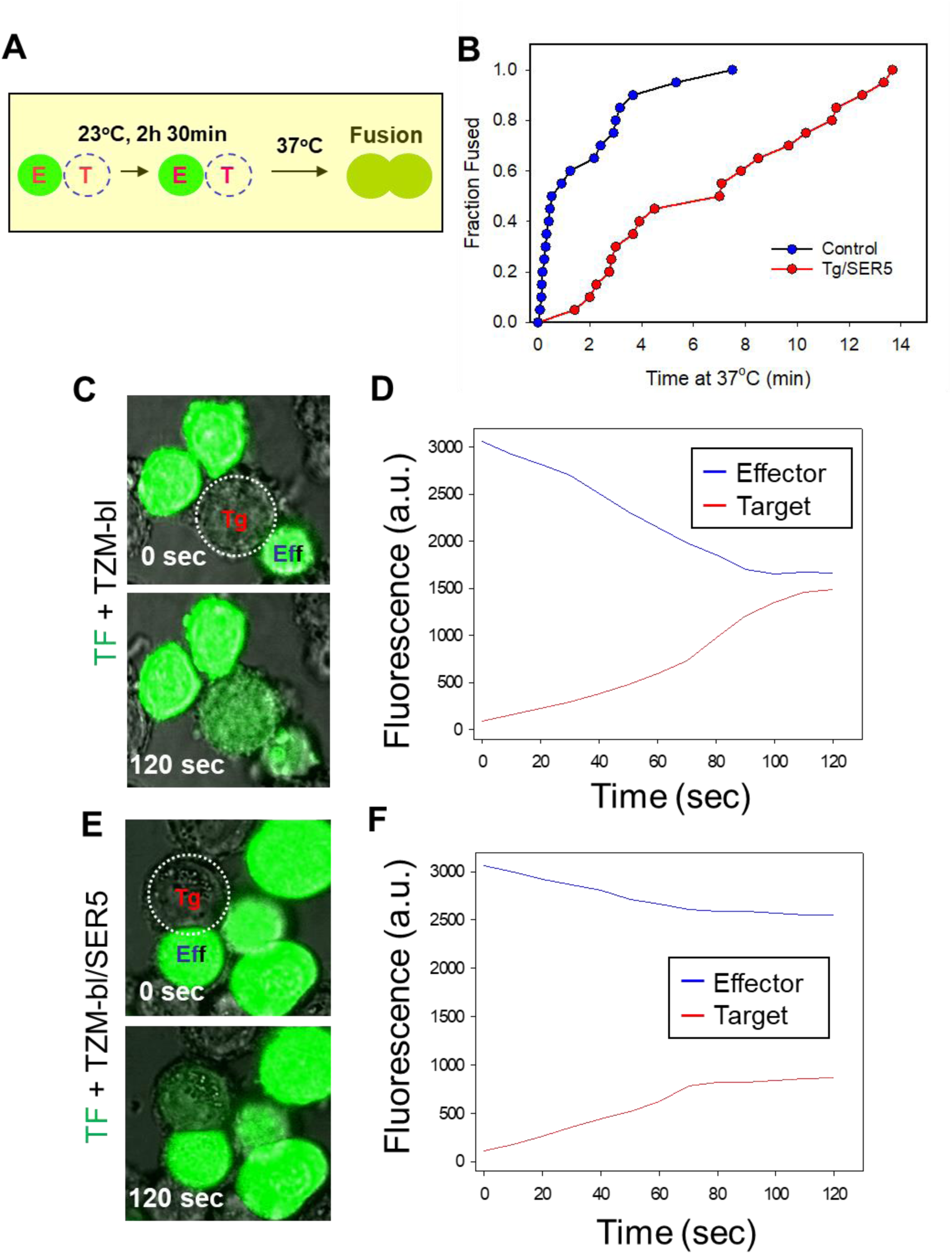
SER5 expression in target cells slows down Env-mediated cell fusion and arrests fusion pore enlargement. The effector TF228.1.16 cells (TF, loaded with calcein AM) and target TZM-bl cells transfected or not with SER5 were co-incubated for 2.5 hours at 23 °C to establish TAS and synchronize cell fusion induced by raising the temperature to 37 °C, using a custom-built temperature jump setup. (A) The kinetics of fusion between individual effector/target cell pairs were monitored by tracking calcein redistribution from effector to target cells following a quick shift to 37 °C. (B, C) Effector TF228.1.16 cells loaded with calcein AM were fused with target TZM-bl cells, using the protocol described in A, and redistribution of calcein between cells was monitored by live cell imaging. Integrated fluorescence intensities of the effector and target cells are plotted in C. (D, E) Same as in panel B, but TF228.1.16 cells were fused with target TZM-bl cells transiently expressing SER5. Integrated fluorescence intensities of the effector and target cells are plotted in E.

We have previously deduced the effective size and dynamics of fusion pores mediated by HIV-1 Env based upon the rate of calcein redistribution between the effector/target cell pairs (29). Strikingly, whereas calcein redistribution from effector to control target cells, was typically completed within 1-2 min, dye redistribution to SER5-expressing cells was much slower and never reached completion (Fig. 5C-F and Suppl. Fig. S7). In other words, none of the fusion events with SER5-expressing target cells allowed calcein to equilibrate between the donor and recipient cell, suggesting a premature closure of small fusion pores. Measurements of the time course and extent of small dye redistribution between a cell pair allows for calculation of a relative pore permeability (29). Plotting multiple pore permeability profiles for either target or effector cells expressing SER5 confirms the virtual lack of pore dilation compared to control cell pairs (Fig. 6A-C). Ensemble averaged profiles of the fusion pore permeability for control and SER5-positive target cells (Fig. 6D) further illustrate this notion. These results demonstrate that SER5 strongly impairs the dilation of Env-mediated fusion pores and, ultimately, promotes pore closure.

**Figure 6.**
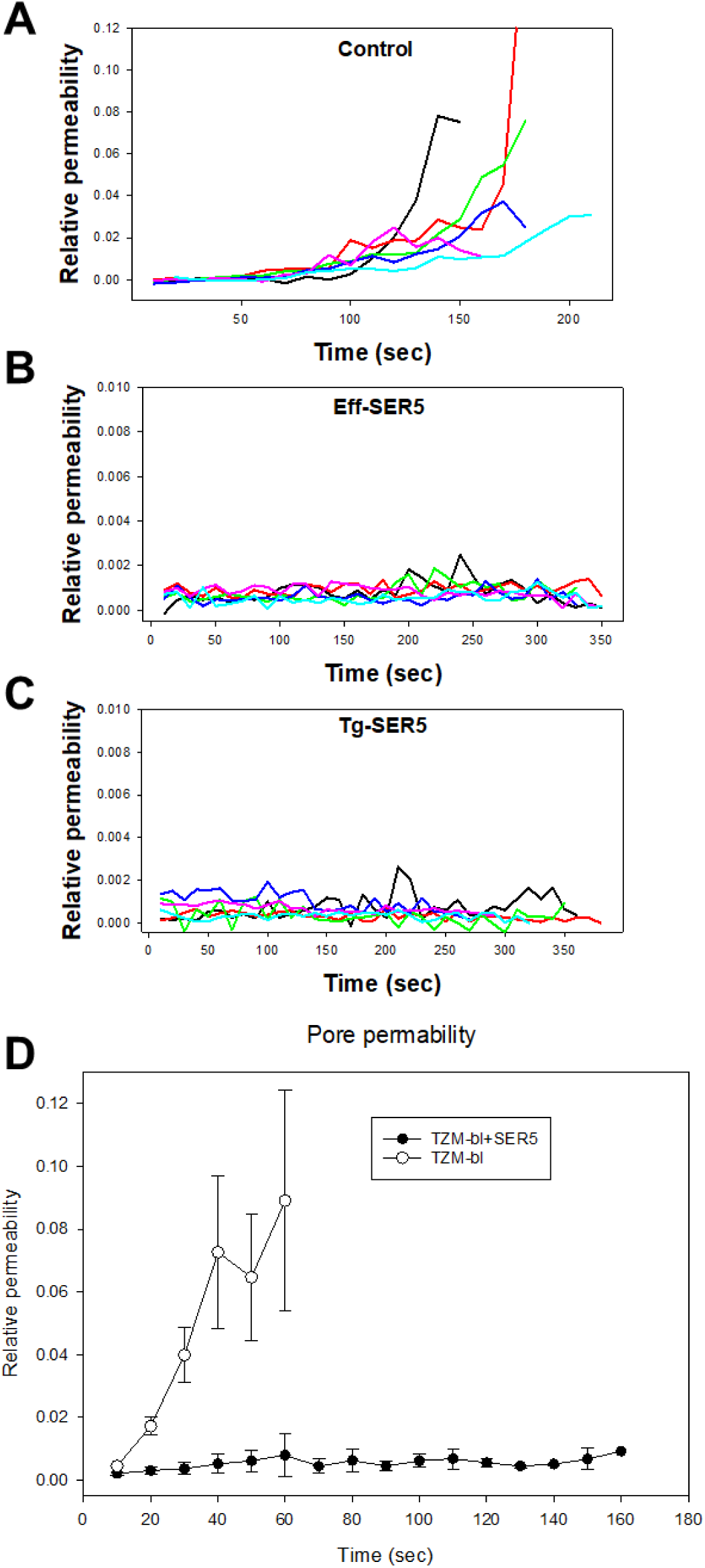
SER5 blocks expansion of fusion pores formed by HIV-1 Env. Fusion pore permeability profiles were calculated from the kinetics of calcein redistribution (exemplified in Fig. 5C-F), as described in Methods. Individual pore permeability profiles between effector TF228.1.16 and target TZM-bl cells for un-transfected TZM-bl cells (A), for SER5-transfected TZM-bl cells (C) and for SER5-transfected TF228.1.16 cells (B) are shown. (D) Average pore growth kinetics were obtained by calculating ensemble averages of multiple pore profiles aligned at the time of pore formation.

### HIV-1 pseudovirus fusion is not inhibited by SER5 expressed in target cells

We next asked if target cells expressing SER5 are less conducive to HIV-1 pseudovirus fusion. Parental TZM-bl cells and cells ectopically expressing SER2-GFP or SER5-GFP (to assess the efficiency of SERINC transfection) were inoculated with pseudoviruses bearing SER5-sensitive (HXB2) Env or –resistant (JRFL) Env, and the resulting fusion was measured using a beta-lactamase (BlaM) assay (53–55). In parallel experiments, we verified that GFP-tagged SER5 retained anti-HIV activity in a cell-cell fusion assay (Suppl. Fig. S8). In stark contrast to robust inhibition of cell-cell fusion by SER5-GFP, neither SER2-GFP nor SER5-GFP expression significantly affected fusion of particles bearing sensitive or resistant Env (Fig. 7A). This result prompted us to test if SER5 in target cells modulates single cycle infection. Although infection of SER5-GFP-expressing cells was modestly reduced compared to control cells, SER2-GFP expression had the same modest inhibitory effect on both HXB2 and JRFL Env-mediated infection (Fig. 7B). This unexpected effect of SER2 on HIV-1 infection, but not fusion, points to a non-specific effect of SERINC expression on post-fusion steps of HIV-1 infection.

**Figure 7.**
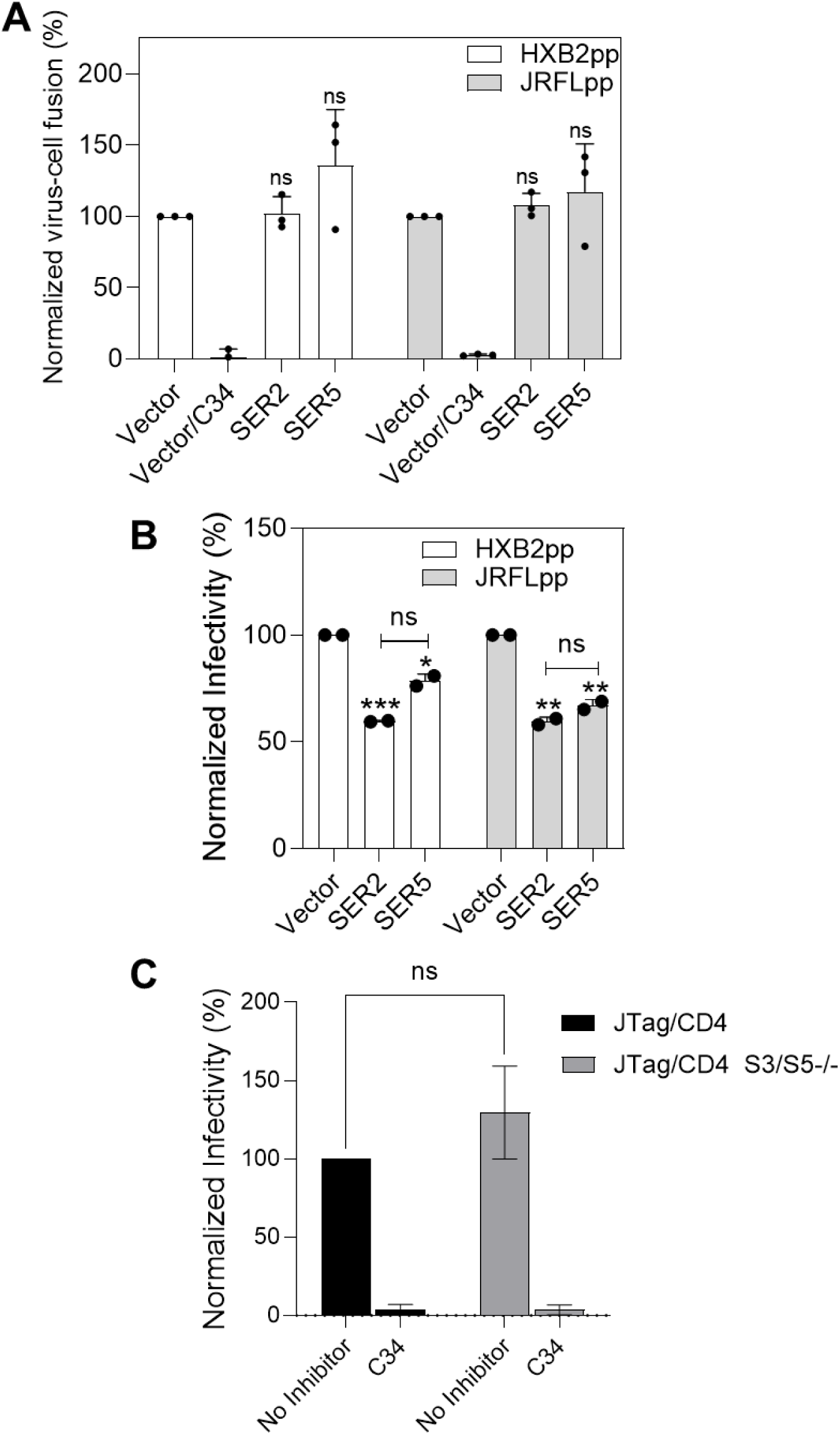
SER5 expressed in target cells does not specifically inhibit HIV-1 pseudotype fusion or infection. The experiments were performed with TZM-bl cells ectopically expressing either SER2-GFP or SER5-GFP (or transduced with an empty retroviral vector). *nef*-negative HIV-1 pseudoviruses harboring BlaM-Vpr and bearing either HXB2 or JRFL Env were used. (A) Virus-cell fusion measured by a BlaM assay. Virus-cell fusion was synchronized by pre-binding the virus in the cold and shifting to 37 °C for 90 min. The C34 inhibitory peptide (1 μM) was added to control wells to block virus fusion. Data are mean and S.D. of three independent experiments, each performed in triplicate. (B) Infectivity of pseudoviruses analyzed in A measured by firefly luciferase activity at 48 h post-infection. Data are mean and S.D. of two independent experiments, each performed in triplicate. (C) The infectivity of *nef*-negative HXB2 pseudoviruses in JTag/CD4 and JTag/CD4 S3/S5−/− cells. In control samples, Env-mediated infection was blocked by C34 peptide (1 µM). Data are mean and S.D. of 4 independent experiments, each in triplicate. Statistical analysis was performed using the student’s *t* test. *, p < 0.05; **, p < 0.01 ***, p < 0.001; ns, not significant.

We further assessed the effect of endogenous SER5 and SER3 expression on HIV-1 entry using parental JTag.CD4 and JTagCD4 S3/S5−/− cells (see Fig. 1A). Consistent with HIV-1 fusion with TZM-bl cells, we did not detect a significant increase in HIV-1 infection of SER3/SER5-negative cells compared to parental cells (Fig. 7C), although 2 out of 4 independent experiments did show a modest enhancement of infectivity (Suppl. Fig. S9).

### SER5 can inhibit HIV-1 pseudovirus-mediated fusion of adjacent cells (fusion-from-without)

To address the discrepancy between the effects of SER5 on cell-cell fusion *vs* virus-cell fusion/infection, we asked whether SER5 can inhibit fusion-from-without (FFWO) – cell-cell fusion mediated by viruses bound to juxtaposed cells (56–58). FFWO involves fusion of two plasma membranes mediated by viral particles fusing with both adjacent cells, without a contribution from internalized virions. We examined FFWO mediated by pseudoviruses displaying SER5-sensitive (HXB2) or -resistant (JRFL) Env, using TZM-bl target cells stably expressing GFP-tagged SER2 or SER5 to readily identified SERINC-expressing cells. SER2-GFP or SER5-GFP expressing TZM-bl cells were non-enzymatically detached from plates, split in two samples that were separately labeled with different cytoplasmic fluorescent dyes. Labeled cells were co-cultured to create a confluent monolayer, followed by virus binding in the cold and incubation at 37 °C to allow FFWO. Microscopic examination of fused (double-positive for both dyes) cells revealed a marked reduction of HXB2 pseudovirus-mediated FFWO of SER5-compared to SER2-expressing target cells (Fig. 8A). By contrast, JRFL pseudovirus-mediated FFWO of SER5-expressing cells was not significantly reduced relative to SER2-expressing cells (Fig. 8A).

**Figure 8.**
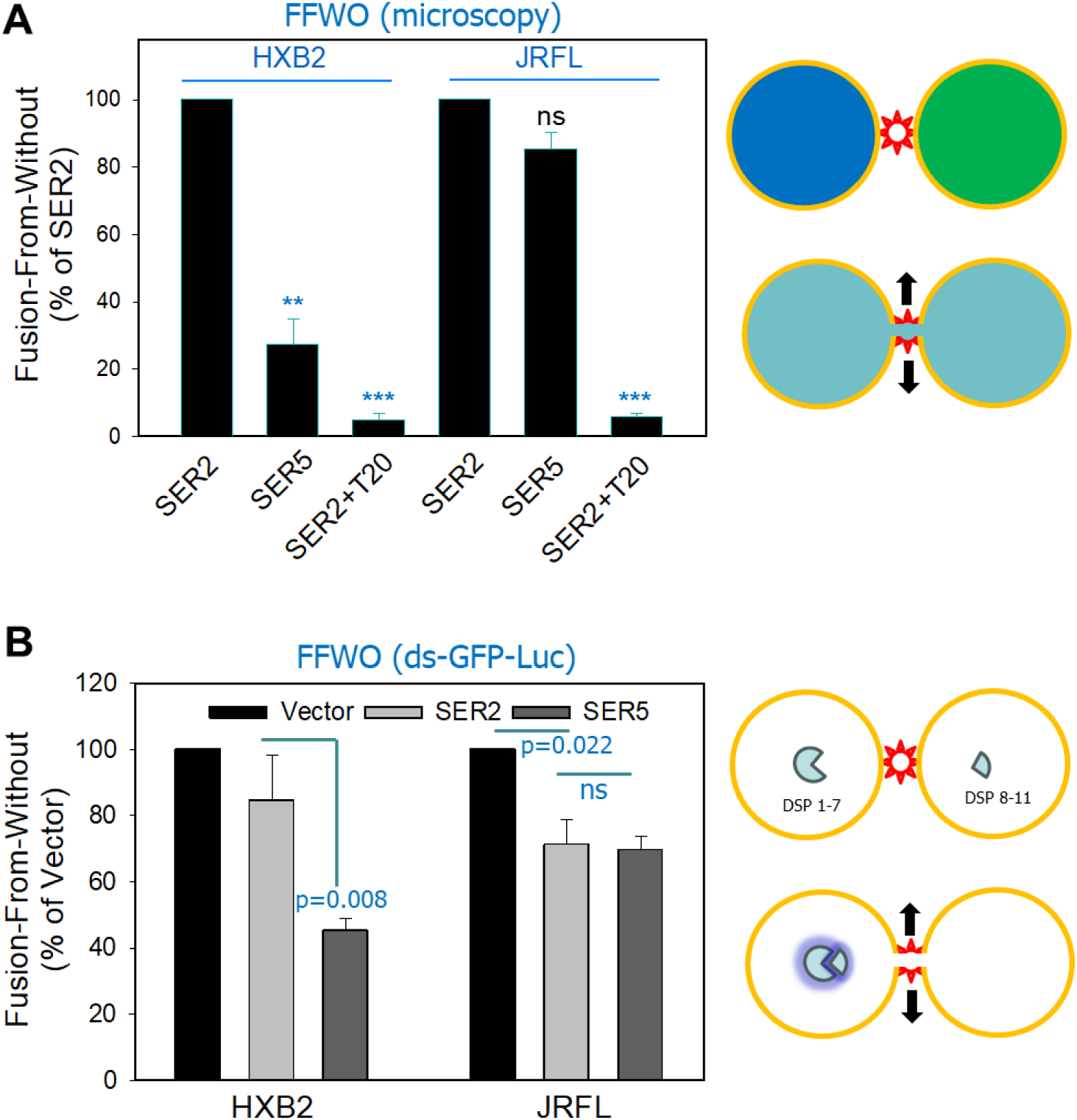
SER5 inhibits HIV-mediated cell fusion (fusion-from-without). (A) TZM-bl cells stably expressing GFP-tagged SER2 or SER5 were used to assess the impact of these proteins on “fusion-from-without” (see the cartoon on the right and Materials and Methods). Control experiments were performed in the presence of the T20 peptide inhibitor (100 nM). Fused (double-positive) cells were quantified microscopically and normalized to the total number of cell-cell contacts. (B) HIV-1 Env-mediated “fusion-from-without” was measured using a dual-split-GFP-luciferase assay (illustrated on the right) using stable TZM-bl cells stably expressing SER2 or SER5 and the complementary fragments of split GFP-Renilla luciferase. Fusion was quantified by measuring the luminescence signal. Results are based on three independent experiments, each done in triplicate, and are expressed as mean ± SEM. Statistical analysis was done using Student’s t-test.

We next verified the effect of SER5 on FFWO using an independent split-luciferase assay. Toward this goal, we used TZM-bl cells stably expressing one of the complementary domains of dual-split GFP-Luciferase (28) and either SER2-GFP or SER5-GFP. HXB2 or JRFL pseudovirus-mediated fusion between these cells results in reconstitution of Luciferase activity. We found that SER5, but not SER2 expression markedly inhibited FFWO mediated by HXB2 particles (Fig. 8B). By comparison, JRFL pseudovirus-mediated FFWO was modestly and equally reduced by both SER2 and SER5 (Fig. 8B). Because SER2 and SER5 expression caused a similar modest reduction in JRFL-mediated FFWO, the effect of SER5 is likely non-specific. Our results thus confirmed that, in contrast to regular HXB2 entry in TZM-bl cells, confining viral fusion to the plasma membrane (FFWO) renders it sensitive to SER5 inhibition.

## Discussion

Here, we demonstrated the ability of SER5 and SER3 expressed in target membranes to inhibit HIV-1 Env-mediated cell-cell fusion and virus-mediated fusion-from-without. This inhibitory effect phenocopied the impact of virus-incorporated SER5 on HIV-1 fusion and infectivity. As with SER5 in the viral membrane, the potency with which SER5-expressed in target cells suppresses fusion is HIV-1 Env strain-dependent (Fig. 1C). Note, however, that a recent study did not observe inhibition of NL4-3 Env mediated cell-cell fusion by SER5 expressed in effector cells (59). Another study also failed to detect an effect of endogenous SER5 levels in HIV-infected Jurkat cells on their fusion with primary macrophages (60). While the reasons for these discrepant results are unclear, we note that both studies examined late fusion products (6-24 hours) that may mask possible kinetic effects of SER5 expression. Also, the second study did not knock out both SER5 and SER3 in Jurkat cells despite the fact that endogenous levels of SER3 in these cells exhibit antiviral activity (5). Neither of the two studies tested the effect of SER5 expressed in target cells.

We also found that SER5 in target or effector cells does not significantly reduce IAV HA-mediated cell-cell fusion (Fig. 2). The lack of SER5 effect on IAV HA-mediated fusion in our experiments agrees with the recent article (14), however, other groups have reported HA subtype- and glycosylation-dependent inhibition of HA-mediated fusion/infection by SER5 expressed on the viral/effector membrane (12–14). SER5 expressed in target cells has been reported to inhibit IAV lipid mixing (hemifusion), apparently, by relocating from the plasma membrane to endosomes upon infection (13) The reasons for these discrepant results are currently unclear but may be related to the levels of SER5 expression.

An important finding of this study is the striking rescue of HIV-1 Env-mediated fusion with SER5-expressing target or effector cells by exogenous PS and PG, but not PC or BMP (Fig. 4). The failure of BMP to antagonize SER5-mediated restriction of Env-mediated fusion indicates that negative charge is not the main determinant of the effect of PS. Of course, we do not know the distribution of exogenously added lipids in cells and, thus, cannot rule out the possibility that a fraction of PS and PG remain at the plasma membrane, while other lipids are removed from the cell surface. Our result in Figure 4 also show a modest increase in fusion efficiency of control cells by exogenous PS and PG, in excellent agreement with our previous finding that cell surface exposure of PS stimulates HIV-1 Env-mediated fusion (51). Considering the recently reported SER5’s lipid scrambling activity (8), we surmise that excess PS may favor a lipid scrambling conformation of SER5, which may be less inhibitory for membrane fusion.

### Possible mechanism of inhibition of Env-mediated fusion by SER5 in target membranes

Several lines of evidence presented in this study support the notion that SER5 exerts its effect on the nascent fusion pores formed by HIV-1 Env, without affecting the upstream steps of fusion. First, SER5 does not interfere with the formation of functional Env-CD4/CXCR4 complexes at a reduced temperature that is not permissive for fusion (Fig. 3). Note, however, that we cannot rule out the possibility that SER5 can slow down CD4 or coreceptor engagement by Env on a time scale that is shorter than that required to establish TAS (>2 hours). Second, SER5 does not capture Env-mediated fusion at a hemifusion stage (Fig. 2). Third, exogenous PS promotes Env-mediated fusion with SER5-expressing cells, regardless of whether it was added to effector or target cells (Fig. 4). This observation is consistent with the SER5’s involvement at the point where two membranes merge and exchange lipids. Finally, we observed a potent destabilization of Env-mediated pores and their collapse upon SER5 expression in either target or effector cells (Figs. 5 and 6). Since SER5 similarly impacts HIV-1 Env-mediated fusion regardless of whether it is present in effector or target cells, we propose that SER5 increases the energetic penalty for sustaining small fusion pores and thereby promotes their collapse.

Inhibition of full HIV-1 fusion with plasma membrane-derived vesicles containing SER3 and SER5 has been observed by cryo-electron tomography (cryo-ET) (61). The authors have reported HIV-1 pseudovirus fusion arrest at hemifusion and early fusion intermediates for viruses containing SER3 or SER5. A major distinction with our results is that these early intermediates were featuring fusion pores comparable to the virus diameter (~100 nm), with fusion products having an hourglass-shaped membrane. In contrast, our definition of small/nascent fusion pores is the ones to limit free dye diffusion and are thus likely in a nanometer range. These types of small pores between HIV-1 and supported membranes that allow the release of mCherry do not appear to be inhibited by SER3 or SER5 relative to lipid mixing (61). By monitoring the fusion pore dynamics in real time, we demonstrate potent destabilization of nascent fusion pores and their collapse caused by SER5 expression in the context of Env-mediated cell-cell fusion. It is possible that these transient events going back to two separate membranes could have been missed by cryo-ET analysis.

Note that the proposed model does not explain the reason for the lack of an effect of SER5 on virus-cell fusion with or infection of TZM-bl or JTag cells (Fig. 7). Surprisingly, SER5 in target cells exhibits distinct effects on HXB2 fusion and on cell-cell fusion mediated by this virus (FFWO, Fig. 8). To reconcile these differences, we propose that these reflect the preferred HIV-1 entry pathway in TZM-bl cells. Our studies have revealed a strong preference for HXB2 entry *via* endocytosis and fusion with pH-neutral intracellular compartments (62). It is thus possible that SER5 effectively restricts HIV-1 Env-mediated fusion at the plasma membrane (cell-cell fusion and FFWO) but fails to interfere with virus entry through an endocytic pathway.

## Materials and Methods

### Plasmids, cell lines and reagents

The expression vectors pCAGGS-HXB2, pCAGGS-JRFL, pcRev, pR9ΔEnvΔNef, pMDG-VSV-G, pBJ5-SER2-HA, pBJ5-SER5-HA, pBJ-SER2-GFP, and pBJ-SER5-GFP have been described previously (4). For experiments in Figure 2B, pcDNA3.1 plasmids expressing HA-tagged SER2, SER3 or SER5 (a kind gift from Dr. S-L Liu, Ohia State University) were used to transfect cells. JR2, AD8, ADA HIV-1 Env expression vectors were previously described (18). Expression vectors for the influenza hemagglutinin strains PR8/34 and X:31were kindly provided by Dr. AL Brass (University of Massachusetts) and Judith White (University of Virginia), respectively. NL4-3.Luc.E-R-(HRP-3418) (63), and pMM310-BlaM-Vpr (ARP-11444) (53), expression plasmids were from the BEI Resources, NIH HIV Reagent Program. The cloning of pQCXIP-SER2-GFP and pQCXIP-SER5-GFP was done as follows. The SER2-GFP fragment was amplified by PCR using TaqDNA polymerase high fidelity (Invitrogen, Waltham, MA, USA, 11304-011), pBJ5-SER2-GFP expression vector as template, and the forward SER2-SbfI 5’-GGCCTGCAGGGCCATGGACGGGAGGATGATGAG-3’, the reverse GFP-NotI 5’-GCTGCGGCCGCTTACTTGTACAGCTCGTCCATGCCGA-3’ primers. The SER5-GFP fragment was amplified using pBJ5-SER5-GFP plasmid as template, and the forward SER5-SbfI 5’-GGCCTGCAGGGCCATGTCAGCTCAGTGCTGTGC-3’ and the same reverse primer as for SER2-GFP. The amplified fragments and the retroviral vector pQCXIP (Clontech, Mountain View, CA, USA, 631516) were digested with SbfI and NotI, purified and ligated. The lentiviral vectors pLenti-DSP1-7 and pLenti6.2-DSP8-11 were a gift from Dr. Z. Matsuda (University of Tokyo) (64).

HEK293T/17 cells were purchased from ATCC (Manassas, VA). HeLa-derived TZM-bl cells (donated by Drs. J.C. Kappes and X. Wu (65) and TF228.1.16, a mammalian Burkitt’s lymphoma cell line that stably expressing HIV-1 envelope glycoprotein (BH-10 clone of HIV-1 LAI, contributed by Drs. Zdenka Jonak and Steve Trulliy (31)) were obtained from BEI Resources. JurkatTag (JTag) and JurkatTag SER3/^−/−^ SER5^−/−^ (abbreviated JTag S3/S5−/−) double-knockout cells were a gift from Dr. H.G. Göttlinger (University of Massachusetts). HeLa cells expressing the HIV-1 ADA Env (HeLa-ADA cells) were a gift from Dr. Marc Alizon (Pasteur Institute, France) (66).

HEK293T/17, HeLa, TZM-bl and COS7 cells were cultured in high-glucose Dulbecco’s Modified Eagle’s Medium (Mediatech, Manassas, VA, USA), while TF228.1.16 cells and JTag-derived cells were grown in RPMI-1640 (Life Technologies, Grand Island, NY, USA). Media were supplemented with 10% FBS (GIBCO BRL) or 10% Cosmic Calf Serum (HyClone, Logan, Utah) and 100 units/ml penicillin/streptomycin (from Hyclone or Gibco). For HEK293T/17 cells the growth medium was additionally supplemented with 0.5 mg/ml G418 (Life Technologies, 1013-027). The JTAg/CD4 versions were obtained by retroviral transduction with pCXbsrCD4 and selection with blasticidin (5). TZM-bl.SER-GFP stable cell lines were obtained by transducing with VSV-G/pQCXIP-SER-GFP pseudotyped viruses and selecting with 1 µg/mL puromycin (InvivoGen, San Diego, CA, USA, ant-pr-1). TZM-bl.SERs-GFP-DSPs stable cell lines were obtained by lentiviral transduction with DSP1-7 or DSP8-11 and selection with 5 µg/mL blasticidin (Research Products International, Mount Prospect, IL, USA, B12200-0.05).

EnduRen^TM^ and Bright-Glo luciferase were from Promega (Madison, WI, USA, E6481), whereas BlaM substrate, CCF4-AM was from Invitrogen (K1089). The HIV-1 gp41-derived C34 and T20 peptides were a gift from Dr. L. Wang (University of Maryland), and from BEI Resources, NIH HIV Reagent Program, respectively. All lipids, DOPC (1,2-dioleoyl-sn-glycero-3-phosphocholine, 850375C), DOPS (1,2-dioleoyl-sn-glycero-3-phospho-L-serine, 840035C), DOPG (1,2-dioleoyl-sn-glycero-3-[phospho-rac-(3-lysyl(1-glycerol))], 840521P), and BMP (bis(monooleoylglycero)phosphate (S,R Isomer), 857133C) were purchased from Avanti Polar Lipids (Alabaster, AL). AMD3100 was from Millipore/Sigma (A5602).

### Pseudovirus production and characterization

All the pseudoviruses were generated by transfecting HEK293T/17 cells grown to ~75% confluency in 10-cm dishes with JetPRIME transfection reagent (Polyplus-transfection, Illkirch-Graffenstaden, France, 114-15). For generating the pseudoviruses used in virus-cell fusion BlaM assay and FFWO, HEK293T/17 cells were transfected with 4 μg pR9ΔEnvΔNef, 2 μg BlaM-Vpr, 0.5 μg pcRev and 3 μg HIV-1 Env expression vector. For luciferase-encoding pseudoviruses used in infectivity assay, HEK293T/17 cells were transfected with 6 μg pNL4-3.Luc.R-E-, 0.5 μg pcRev and 3 μg HIV-1 env expression vector. The viral supernatants were collected at 40 - 48 h post-transfection, filtered through 0.45 μm polyethersulfone filters (PES, VWR), and concentrated 10× using Lenti-X concentrator (Takara, San Jose, CA, USA, 631232,). The p24 content of pseudoviruses was determined by ELISA [doi: 10.1016/j.vaccine.2007.09.016], and the infectious titer was determined by β-Gal assay in TZM-bl cells (67).

### Virus-cell fusion, fusion from without (FFWO) and infection assays

The virus-cell fusion using a β-lactamase (BlaM) assay was done, as previously described (67). Briefly, HIV-1 pseudovirus (MOI ~1) bearing respective envelope glycoprotein and β-lactamase fused to Vpr (BlaM-Vpr) was bound to a confluent monolayer of target cells in 96-well black clear bottom by centrifugation at 4 °C for 30 min at 1550×g. The unbound virus was washed out, growth medium was added, and samples were incubated at 37 °C, 5% CO_2_ for 90 min, after which the cells were loaded with CCF4-AM substrate and incubated at 12 °C overnight. The blue to green fluorescence ratio was measured using the Synergy HT fluorescence microplate reader (Agilent Bio-Tek, Santa Clara, CA, USA).

For FFWO experiments, two different assays were used: (i) a microscopy-based assay using cytosolic dyes, and (ii) a luciferase assay using DSPs expressing cells. For microscopic detection of FFWO, TZM-bl cells stably expressing SER2-GFP or SER5-GFP, each preloaded with either a blue dye CMAC (10 µM, ThermoFisher, C2110) or 2 µM Calcein Red-Orange AM (C34851) for 30 min at 37 °C. A 1:1 mixture of cells loaded with CMAC or Calcein Red-Orange were co-cultured overnight. Next day, HIV-1 pseudoviruses (MOI ~4) was added to confluent cultures and spinoculated at 10 °C for 30 minutes, 1500 x g. Cells were washed and incubated for 2.5 hours at 37 °C. Fused (double-positive) cells were quantified microscopically and normalized to the total number of cells. For luciferase-based detection, TZM-bl.SER-GFP.DSP1-7 and TZM-bl.SER-GFP.DSP8-11 were seeded at a 1:1 ratio in 96-well black-clear bottom the day before the experiment. On the day of the experiment, cells were loaded with 60 μM EnduRen™ substrate for 1 hour at 37 °C, 5% CO_2_. Next, HIV-1 pseudoviruses (MOI~4) was added to cells and spinoculated at 10 °C for 30 minutes, 1500 x g. After washing out the unbound virus, fresh growth medium was added, and cells were incubated for 3 hours at 37 °C. The luciferase signal was measured using a Wallac1420 multilabel counter (PerkinElmer, Turku, Finland).

For infectivity assay, 1·10^5^ JTag/CD4 or JTag/CD4 S3/S5−/− cells were inoculated with 2 IU/cell HXB2 pseudoviruses bearing luciferase by centrifugation at 4 °C for 30 min at 1550×g, and incubated at 37 °C, 5% CO_2_ for 48 h. Cells were lysed using Bright-Glo luciferase, and luciferase signal was measured on a TopCount NXT reader (PerkinElmer Life Sciences, Shelton, CT, USA)

### Cell-Cell Fusion

Cell-cell fusion was measured, as previously described (29, 30). Briefly, effector cells were loaded with Calcein AM (ThermoFisher, C3099) and target cells were loaded with CMAC Blue (ThermoFisher, C2110). The cells were detached from plates using PBS supplemented with EDTA and EGTA, resuspended in PBS++, mixed at 1:1 ratio, and allowed to attach to poly-L-lysine–coated 8-well chamber slides (Millipore/Sigma, P1274; Lab-Tek, 177402) for 30 min at 23 °C. Fusion was triggered by applying an acidic (pH 5.0) buffer at 37 °C. Where indicated, cells were treated for 1 min at room temperature with chlorpromazine (CPZ; Millipore/Sigma, 215921).

### Monitoring fusion pore enlargement

Effector TF228.1.16 cells were loaded with calcein-AM, mixed with target TZM-bl cells, and allowed to adhere to poly-L-lysine–coated coverslips for 30 min at room temperature. Fragments of coverslips containing cells were transferred to a custom-built imaging chamber and placed onto an IR-absorbing coverslip at the bottom. Local temperature was rapidly raised and maintained at 37 °C by illuminating the absorbing coverslip with an IR diode, as described in (29). Dye transfer was monitored using a Fluoview 300 laser-scanning confocal microscope (Olympus IX70, Melville, NY, USA) with a UPlanApo 60×/1.20 NA water-immersion objective, and pore permeability as a function of time was calculated, as described previously (29).

### Flow Cytometry

Cells (~10^6^) transfected or not with SER2 or SER5 were immunostained with either the human anti-CD4 antibody (SIM2, #723) or the mouse anti-CXCR4 antibody (12G5, #3439) both obtained from the NIH HIV Reference Reagents Program. Antibodies were diluted 1:100 in PBS++ supplemented with 5% goat serum and 0.1% sodium azide, for 45 min on ice. After washing with the same buffer, cells were incubated with the appropriate secondary antibody FITC-conjugated goat anti-human IgG (Millipore/Sigma, F9887) or Alexa Fluor 488–conjugated goat anti-mouse IgG (ThermoFisher, A10667) diluted 1:500 for 45 min on ice. Following multiple washes, cells were resuspended in 800 μL PBS-- and analyzed by flow cytometry using a Guava EasyCyte System (Hayward, CA).

### Statistical analysis

Unpaired Student’s t-test or Mann-Whitney test were done either using GraphPad Prism version 9.3.1 for Windows (GraphPad Software, La Jolla, CA, USA), or SigmaPlot version 15.0 (Systat Software, Inc., Palo Alto, CA).

## Acknowledgments

The authors wish to thank Hui Wu for excellent technical support. This work was supported by the NIH R37 AI150453 grant to GBM.

